# Towards Improving Embryo Selection: Simultaneous Next Generation Sequencing Of DNA And RNA From A Single Trophectoderm Biopsy

**DOI:** 10.1101/277103

**Authors:** Noga Fuchs Weizman, Brandon A. Wyse, Ran Antes, Zenon Ibarrientos, Mugundhine Sangaralingam, Gelareh Motamedi, Valeriy Kuznyetsov, Svetlana Madjunkova, Clifford L. Librach

**Affiliations:** CReATe Fertility Centre, Toronto, Canada; Department of Obstetrics and Gynecology, University of Toronto, Toronto, ON, Canada; Department of Physiology, University of Toronto, Toronto, ON, Canada; Department of Gynecology, Women’s College Hospital, Toronto, ON, Canada

**Author notes:** Authors contributed equally (Noga Fuchs Weizman, Brandon A. Wyse). **CORRESPONDING AUTHOR:** Dr. Clifford L. Librach, 790 Bay St. – Suite 1100 Toronto, ON, Canada; or Dr. Svetlana Madjunkova, 790 Bay St. – Suite 1100 Toronto, ON, Canada.

**Keywords:** Next generation sequencing, RNA-sequencing, embryo selection, preimplantation genetic assessment, multi-omic analysis

## Abstract

Improved embryo selection is crucial in optimizing the results from assisted reproduction. Preimplantation genetic screening reduces time to pregnancy and miscarriages. Correlating the transcriptome of an embryo, with fertility treatments and outcomes, holds promise in improving the overall results. We developed a novel method for embryo selection in fertility treatments that integrates embryonic genomic and transcriptomic data and evaluated it in this pilot study.

A total of 21 embryos donated for research were included. Three were used for the initial development and optimization of sample processing and sequencing. Thereafter, 18 embryos were used to demonstrate the clinical safety and reproducibility of our method. Two trophectoderm biopsies were taken from each embryo: one was processed as a clinical sample for genomic profiling (control, n=18), while the other biopsy (n=18) was split and utilized for independent, simultaneous genomic and transcriptomic analysis, here termed Preimplantation Genetic and Transcriptomic Testing (PGT^2^).

High quality genomic and transcriptomic data were obtained from all analyzed samples. The concordance between genomic data obtained with PGT^2^ and control samples was 100% with clinical grade quality metrics. Euploid embryos showed downregulation of genes involved in anaerobic metabolism, oxidative phosphorylation, and fatty-acid oxidation. This is the first study to provide full genomic and transcriptomic profiles from a single TE biopsy from human embryos in a clinical setting unleashing the potential of improving embryo selection and outcomes in infertility treatments. Clinical trials are needed to correlate transcriptomic data with outcomes.

**SUMMARY:** Despite advances in assisted reproductive technologies, the success rate has remained relatively constant. Under the age of 35, there is a 40% chance of delivering a child per embryo transfer, which decreases with increasing maternal age. Prioritizing embryos for transfer is based on morphological assessment and, in some cases, incorporates genetic testing as well. Selection of euploid embryos for transfer shortens the time to pregnancy and reduces the risk for miscarriages. Adding the mRNA analysis to the genomic assessment of an embryo has the potential of improving the outcomes of fertility treatments.

## INTRODUCTION

With elective single embryo transfer (eSET) becoming the leading strategy to avoid multiple gestations in assisted reproductive technologies (ART), improved embryo selection is crucial to maximize pregnancy rates per embryo transfer. Morphological and/or morphokinetic criteria have limited predictive value for implantation potential and ongoing pregnancy rates (JONES *et al.* 2008; KASER AND RACOWSKY 2014). Preimplantation genetic testing for chromosomal aberrations (PGT-A) reduces time to pregnancy and helps avoid miscarriages, but it cannot predict the implantation potential of a euploid embryo (DAHDOUH *et al.* 2015).

Implantation failure is believed to be due to a range of factors including chromosomal abnormalities, asynchrony between embryo development and uterine receptivity and factors associated with treatment interventions and techniques (JONES *et al.* 2008). Several molecular processes and pathways involved in implantation have been characterized (WELLS *et al.* 2005). However, the complete molecular dialogue between the maternal and the fetal components leading to implantation, in particular the role of the blastocyst, still remains poorly understood (MANTIKOU *et al.* 2016). Defining what differs between viable and nonviable embryos and specifically, understanding the process of implantation is important for optimizing infertility treatment outcomes for all the infertile population (KIRKEGAARD *et al.* 2015).

Previous studies focusing on defining implantation potential of blastocysts suffer from several limitations. Most available knowledge of the molecular basis of preimplantation embryo development comes from gene expression studies on mouse, bovine and non-human primate embryos (HAMATANI *et al.* 2006; WANG AND DEY 2006; GUPTA *et al.* 2017; NAKAMURA *et al.* 2017; SALEHI *et al.* 2017). Earlier studies looking at cellular pathways that are activated during early embryonic stages and may be necessary for successful implantation have done so via RT- PCR (MARIN *et al.* 2017). The pathways that were studied include cell cycle regulation, DNA repair, apoptosis, and maintenance of accurate chromosomal segregation and construction of the cytoskeleton (WELLS *et al.* 2005; ASSOU *et al.* 2011; MARIN *et al.* 2017). Basing such investigations on RT-PCR is highly selective, focusing on specific genes or pathways and can introduce significant bias to the data. Moreover, most of the available data comes from studies using pooled samples or from functional models of human embryonic cells that cannot accurately reflect the potential of an individual sample (PARKS *et al.* 2011). This impedes our ability to predict outcome owing to variability that exists between individual blastocysts.

Yan et al 2013 were the first to perform transcriptome analysis by Next Generation Sequencing (NGS) on single cells derived from human embryos at different developmental stages, thereby better defining the pathways involved in early embryonal development (YAN *et al.* 2013). Followed by Petropoulos et al 2016, that extensively studied the lineage segregation at different stages of development (PETROPOULOS *et al.* 2016). Recently, several studies identified genes that were differentially expressed between blastocysts resulting in healthy fetal development and blastocysts that failed to implant (EL-SAYED *et al.* 2006; JONES *et al.* 2008; PARKS *et al.* 2011; KIRKEGAARD *et al.* 2015). Notably, these studies did not control for the ploidy status of the tested embryos, thereby affecting our ability to know whether differentially expressed genes are affected by the ploidy or whether they represent a true finding (IQBAL *et al.* 2014).

There is still an unmet need for methods to define developmental and implantation potential of a given embryo, alongside its ploidy status. The ideal embryo selection tool should be clinically applicable, synergistic with currently available methods, and improve outcomes. Our aim was to create a clinical tool for simultaneous assessment of chromosomal copy number by low pass whole genome sequencing, and transcriptomic profile using whole transcriptome RNAseq. Being able to provide genomic and transcriptomic data from a single biopsy sample comprised of 4-6 trophectoderm cells is another step on our way to a multi-omic assessment of embryos.

## MATERIALS AND METHODS

### Blastocyst selection, collection, and biopsy

This research received approval from the University of Toronto Research Ethics Board (#30251). A total of 20 blastocysts donated for research from a total of 16 patients were used for this study. Two fresh blastocysts deemed not suitable for transfer were used for cell lysis and sequencing library optimization. Eighteen additional frozen blastocysts that were previously biopsied for genetic testing and deemed unsuitable for transfer due to positive findings for single gene disorders or chromosomal aneuploidy were also utilized for the study: 7 euploid blastocysts were donated for research after a positive preimplantation genetic diagnosis of a known genetic mutation along with 11 aneuploid blastocysts that were diagnosed by PGT-A. These blastocysts were thawed, cultured until re-expanded and their trophectoderm (TE) was re-biopsied (4-6 cells) by one of our clinical embryologists following the standard clinical operating procedures for TE biopsy. Figure S1 contains representative images of blastocysts and embryo biopsies. From each embryo two separate TE biopsies were performed where one TE biopsy was processed as a clinical sample and the other TE biopsy was utilized for evaluation of the new method. For the purpose of transcriptomic data analysis, we grouped these samples into 2 cohorts: one cohort included samples from embryos that were expected to implant and survive the first trimester, whereas the other cohort consisted of samples expected to either not implant or implant and not survive through the first trimester (12 gestational weeks) of pregnancy.

### Patient Demographics

Patient demographic information such as age, BMI, AMH levels, stimulation information and outcomes was collected from the patients’ clinical charts and that data was anonymized by assigning a non-identifying study number. Patient demographics are presented in Table 1.

### Optimization of cell lysis method for TE biopsy

Two blastocysts were used for optimizing the cell lysis step and DNA/cDNA library preparation for NGS. Figure 1 depicts the detailed workflow of cell lysis optimization.

**Figure 1:**
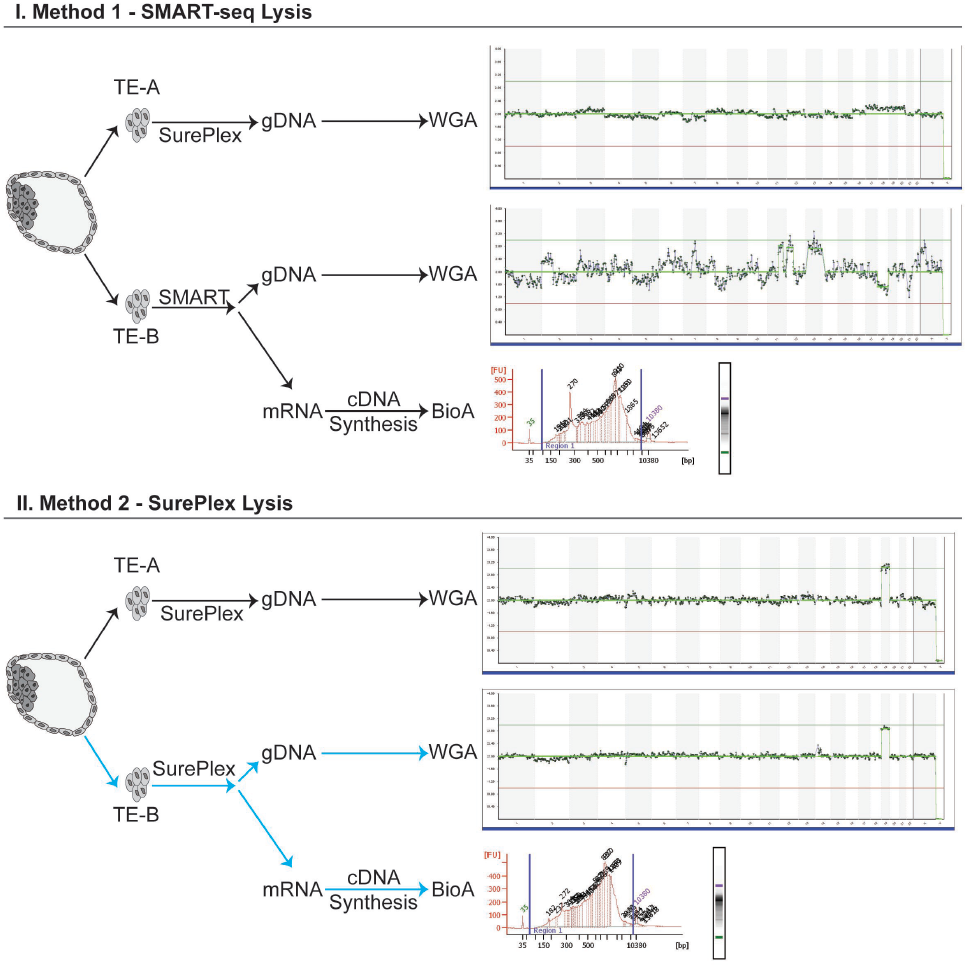
Detailed workflow of the cell lysis optimization to obtain both high quality gDNA and mRNA from the same TE biopsy. TE-A is the “clinically representative” control biopsy, lysed and processed using the standard clinical workflow for PGT-A. TE-B is the test biopsy, where cells are lysed with either SMART (Method 1) or SurePlex (Method 2) kits. The lysate was split and processed according to the standard SurePlex protocol for gDNA amplification, or the standard SMARTseq protocol for cDNA synthesis. Lysis of biopsied cells with SurePlex yields high quality gDNA and mRNA. From each blastocyst cDNA was synthesized, amplified, and its integrity/quality was assessed by BioAnalyzer 2100 (Agilent Technologies, CA). All samples, regardless of lysis method, produced high quality cDNA. However, only the sample lysed using SurePlex (Method 2) produced both high quality cDNA and gDNA which passed all clinical quality control metrics after NGS using VeriSeq Kit (highlighted with blue arrows).

From each of the embryos, we obtained two separate TE biopsy samples (4-6 cells) (Figure 1). The first TE biopsy sample (TE-A) from each embryo was processed as a “clinical sample” and used as a control. TE-A was lysed and amplified using the SurePlex WGA kit (Illumina, CA), which is the current standard of practice in preimplantation genetic testing for chromosomal aberrations (PGT-A). The second TE biopsy sample (TE-B) was lysed using one of the two different lysis methods: I. SMART-seq (Takara BioInc, CA) or II. SurePlex kit (Illumina, CA). In lysis option I the biopsied cells were deposited into 2.5ul of DNase/RNase free water and lysed using SMART-seq lysis mix. For lysis option II, biopsied cells were deposited into 2.5ul of 1xPBS buffer and lysed using cell extraction enzyme mix from SurePlex WGA kit. The lysates were subsequently split and independently processed for both gDNA WGA and cDNA synthesis. The resulting WGA-gDNA was sequenced using Nextera XT library preparation for NGS following the VeriSeq protocol, described in detail below (Illumina, CA). cDNA quality was assessed using BioAnalyzer 2100 (Agilent Technologies, CA). Table 2 presents the results for chromosomal aberrations and sequencing quality metrics from VeriSeq sequencing of gDNA obtained through SurePlex WGA kit using lysis with I. SMART-seq (Takara BioInc, CA) or II. SurePlex kit (Illumina, CA).

### Whole genome amplification, gDNA sequencing, and analysis

Whole genome copy number variation (CNV) analysis was done by whole genome low pass (0.1x) NGS, using the VeriSeq PGS Kit (Illumina, CA). Briefly, after whole genome amplification (WGA) was performed, according to manufacturer’s instructions, gDNA was tagmented and amplified. The amplified DNA was indexed, purified using AMPure XP beads (1:1 ratio), and normalized using magnetic beads. The normalized libraries were pooled, denatured, and sequenced using a MiSeq (single-end, 1× 36bp). BlueFuse Multi (Illumina, CA) was used for chromosome CNV analysis and data visualization. The optimal metrics for PGT-A are 500,000 reads passing filter and a sample noise score (Derivative Log Ratio - DLR) of <0.2; 250,000 reads and DLR <0.4 is clinically acceptable.

### Accuracy of the PGT-A results using PGT^2^ method

To ensure that splitting the cell lysate for simultaneous sequencing of gDNA and RNA does not impact the clinical PGT-A result, we obtained clinical grade TE biopsies from 18 previously frozen embryos with a known ploidy status. In each of these cases the TE biopsy was lysed and then split: half the lysate underwent WGA and VeriSeq sequencing and the other half of the lysate underwent cDNA synthesis and RNA sequencing (Table 3). cDNA quality and quantity were confirmed using a BioAnalyzer 2100 (Agilent Technologies, CA). The WGA product quantity and quality were confirmed by fluorimeter (Qubit - ThermoFisher) and electrophoresis (2% w/v agarose, 100V for 30min), respectively.

### Low-input cDNA synthesis and amplification

cDNA was synthesized and amplified using the SMART-Seq v4 Ultra Low Input RNA Kit for Sequencing (Takara Biosciences, CA) according to the manufacturer’s instructions. Briefly, biopsied cells were deposited into DNase/RNase free water, lysed using the optimized SurePlex lysis method, and mRNA was selectively reverse transcribed using an oligo-dT primer (3’ SMART-Seq CDS Primer II A). The single-stranded cDNA was amplified and purified using AMPure XP beads (Beckman Coulter, MA). cDNA size distribution was assessed using BioAnalyzer 2100 high sensitivity DNA chip (Agilent Technologies, CA).

### cDNA library preparation and RNA sequencing

One nanogram of cDNA was used as input into Nextera XT (Illumina, CA) and libraries were generated according to manufacturer’s instructions. Briefly, cDNA was tagmented, amplified, and indexed. The indexed libraries were purified using AMPure XP beads (1:1 ratio) (Beckman Coulter, MA). Libraries were quantified by Qubit and 2100 BioAnalyzer and normalized using the KAPA Library Quantification kit (Roche, CA). Normalized libraries were pooled, denatured, and 1.2pM was loaded onto a NextSeq High Output (300 cycle) flowcell and sequenced (paired-end, 2x127bp) on NextSeq550 (Illumina, CA).

### RNAseq bioinformatic analysis

Reads were trimmed based on read quality (Phred>28) and aligned and quantified to hg19 using STAR (Spliced Transcripts Alignment to a Reference). Low abundant transcripts were excluded (maximum<20 in 80% of samples) and normalized using Trimmed Mean of M-values (TMM). Normalized read counts were further filtered (excluded if maximum<5 in 50% of samples) and used to perform principal component analysis (PCA) clustering and hierarchical clustering to elucidate the inter-sample variability between the samples. We conducted differential expression (DE) using DESeq2 (LOVE *et al.* 2014) comparing euploid to all aneuploid embryos, and monosomy 16 (M16) to trisomy 16 (T16) embryos. Significantly differentially expressed genes were defined as FDR<0.05 and fold change (FC) < -2 or FC>2. Gene Set Enrichment Analysis (GSEA) was conducted to determine what gene pathways/gene sets are impacted by the gain or loss of chromosome 16 when compared to euploid blastocysts. This analysis was conducted in Partek Flow (version 7.0.18.0218, MO) and the pipeline is available upon request.

### qPCR Validation of RNAseq data

Four TE lineage specific gene markers were chosen based on previous literature (KIRKEGAARD *et al.* 2015; PETROPOULOS *et al.* 2016). Validated gene-specific probe based PrimeTime™ qPCR assays (IDT, IL) were used for validation of RNA-seq NGS results with the housekeeping gene, *RPLP0* as the reference gene. Each PrimeTime™ gene-specific assay consists of two exon-spanning primers and a gene-specific fluorogenic probe labeled with FAM, and an internal and terminal non-fluorescent quencher (ZEN and Iowa Black FQ, respectively). All selected targets were assayed in duplicate using PrimeTime™Gene Expression Master Mix (IDT, IL) using the following cycling conditions: polymerase activation at 95°C for 3 min; 45 cycles of 15 s denaturation at 95°C and 1 min annealing/extension at 60°C. Relative fold change (ΔΔCt) method was employed to quantify gene expression. Data Analysis was performed using GraphPad Prism (version 5.02). The list of primers and probes used for validation is given in Table S1.

### Data Availability Statement

The bioinformatic pipeline is available upon request as are the complete list of gene sets from pathway analysis using GSEA. The authors affirm that all data necessary for confirming the conclusions of the article are present within the article, figures, and tables.

## RESULTS

### Blastocyst collection and biopsy

Twenty blastocysts were biopsied for this study; 2 were used for lysis and sequencing library optimization, and 18 were used for simultaneous sequencing of gDNA and RNA. Tables 2 and 3 present details on the analyzed embryos, biopsies taken, and methods used.

### Optimization of TE biopsy cell lysis

As described in Figure 1 we compared the performance of two different cell lysis methods (I. SMART-seq (Takara BioInc, CA) or II. SurePlex kit (Illumina, CA)). in obtaining gDNA and RNA from the same sample. Biopsied cells lysed using the SMART-seq protocol (I) (Takara BioInc, CA) yielded cDNA of high quality and quantity, measured by BioAnalyzer 2100 (Agilent Technologies, CA) and fluorometer, respectively. The quantity of gDNA from this lysis approach was in the expected range (Table 2) however, the integrity was affected which resulted in poor low pass whole genome NGS results using VeriSeq kit (1,092,118 reads, DLR 0.43) (Figure 1). The SurePlex lysis method (II) yielded high quality and quantity of cDNA and gDNA yielding high quality NGS results using VeriSeq kit (638,468 reads, DLR 0.16). Based on these findings, we established our novel PGT^2^ method: 1. Single TE biopsy, 2. SurePlex cell lysis mix, 3. lysate splitting and 4. simultaneous independent gDNA and cDNA synthesis (Indicated in Figure 1 with blue arrows).

### Concordance of PGT-A results between standard PGT and the PGT^2^ methods

Table 3 summarizes the clinical preimplantation genetic testing results for chromosomal aberrations and results obtained from additional TE biopsies using the PGT^2^ method from 18 embryos donated for research.

There was a 100% concordance between clinical PGT-A results and PGT^2^ from the same blastocysts using the same VeriSeq PGS kit were concordant with the chromosomal CNV analysis by NGS of the gDNA from the split biopsy in all 18 blastocysts (Table 3). All samples produced sufficient quantity and high-quality cDNA for RNAseq by BioAnalyzer 2100 (Agilent Technologies, CA) (Figure S2).

### The TE transcriptome is dramatically altered by ploidy

When comparing euploid samples to samples with monosomy or trisomy 16 (-16 or +16) using Principal Component Analysis, this transformation is defined so that the first PC explains the largest amount of variability in the dataset. Using this analysis, 77.1% of the variability is explained by PC 1, 2, and 3 (Figure 2). Along the first PC (PC1=63.2%), a tight cluster of euploid biopsies is separated from monosomy or trisomy 16 (T16) samples. Interestingly, T16 and monosomy 16 (M16) samples diverge along the second PC (PC2=7.5%). Hierarchical clustering based on normalized read counts further demonstrates the significant differences between euploid and M16 or T16 and highlights samples and genes of interest (Figure 2D).

**Figure 2:**
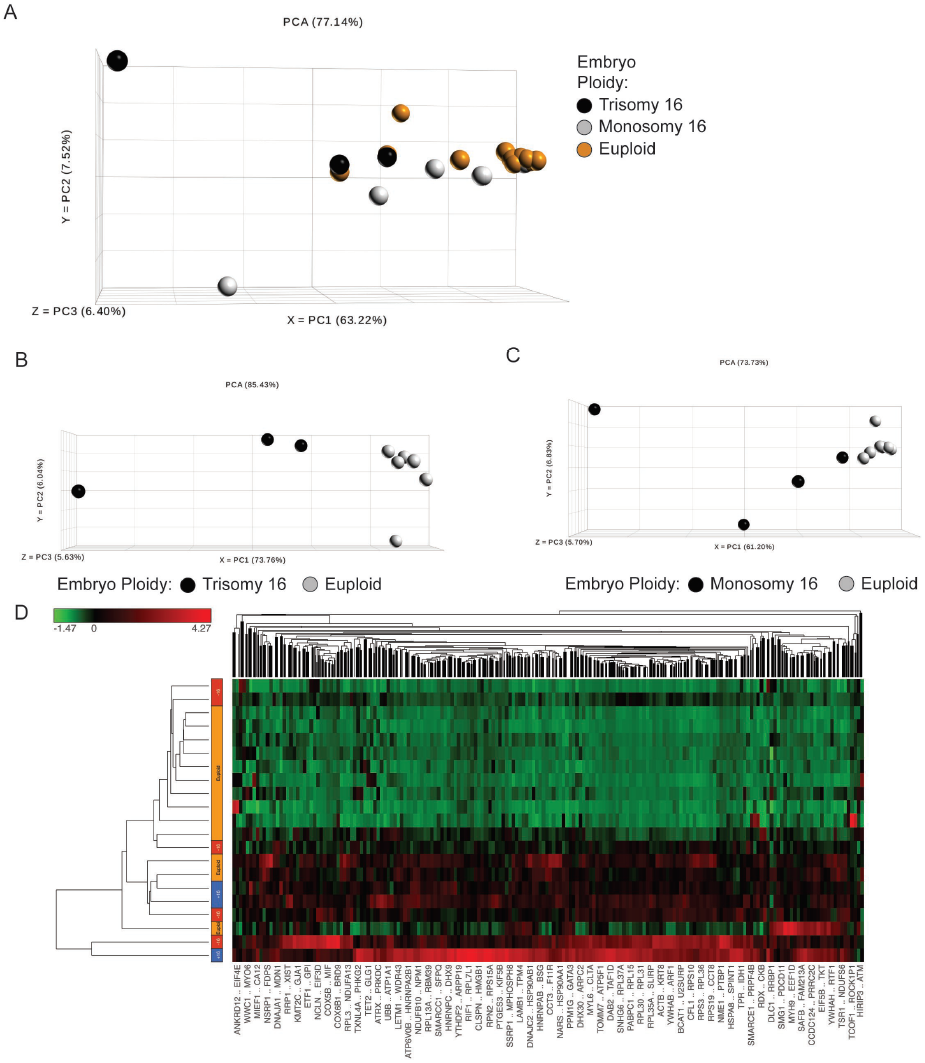
Blastocyst ploidy dramatically alters the trophectoderm transcriptome. A) Principal component analysis of all blastocysts (trisomy/monosomy 16, n=8 and euploid, n=7) shows significant separation along PC1 and PC2 by ploidy. This is further exaggerated when comparing trisomy (n=3) or monosomy 16 (n=5) alone to euploid (n=7), (B) and (C), respectively. D) Samples also cluster under unsupervised hierarchical clustering when plotting standardized expression values. Here, samples are on the rows and features on the columns (red indicating high standardized expression, green indicating low standardized expression).

### RNA transcripts differ between euploid embryos and embryos with a single aneuploidy

Several genes that are active in placentation were differentially expressed between euploid and aneuploid embryos including *FADS1*(-8.9 fold), *PGK1*(-7.7 fold), and *PPAT* (-11.7 fold). When comparing euploid embryos to either monosomy or trisomy 16 embryos; genes associated with mitochondrial respiration (i.e. *STOML2*), angiogenesis (i.e. *EMP2*), and cell migration and invasion (i.e. *PTTG1*) were differentially expressed. Several genes involved in cell cycle regulation, DNA damage repair (*RHOBTB1* and *DDX19B, GDF15*), and cell signaling (*FGFR2*) were differentially expressed between embryos with T16 and M16. All differentially expressed genes across different comparisons are presented in Figure 3 and Table S2-5.

**Figure 3:**
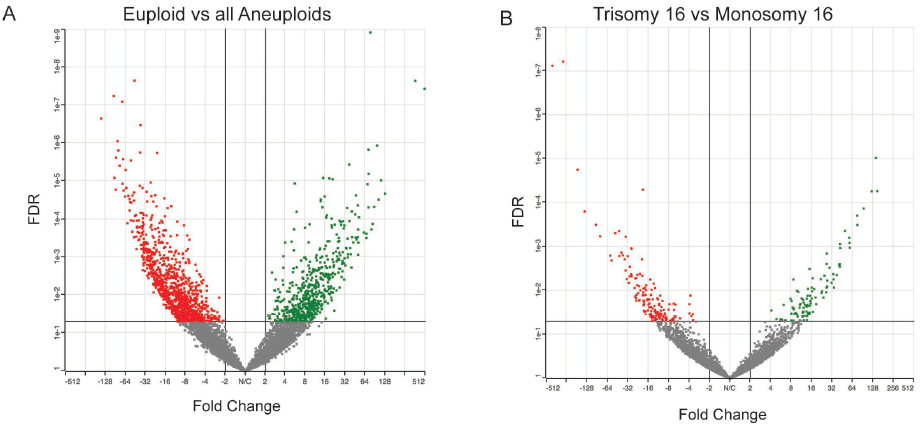
Differential expression analysis between euploid and aneuploid blastocysts. A) When comparing euploid vs all aneuploid biopsies using DEseq2, 1574 transcripts were differentially expressed (1001 down-regulated and 573 up-regulated). B) When comparing trisomy 16 vs monosomy 16 using DEseq2, 251 transcripts were differentially expressed (155 down-regulated and 96 up-regulated). Red indicates significantly downregulated (FC<-2) and green significantly upregulated genes (FC>2) at FDR<0.05.

### Euploid embryos significantly downregulate pathways involved in energy metabolism, transcription, and translation

When comparing TE samples from euploid embryos with TE samples from aneuploid embryos, 156 gene sets were enriched for downregulated genes in the euploid cohort (FDR<0.1) (data not shown). Most notably, pathways involved in energy metabolism (fatty acid oxidation, gluconeogenesis, and mitochondrial translation termination) were significantly enriched in downregulated genes in the euploid cohort (Figure 4). Moreover, pathways involved in nucleotide synthesis, RNA transcription and protein translation were also enriched in downregulated genes in the euploid cohort (Figure 4).

**Figure 4:**
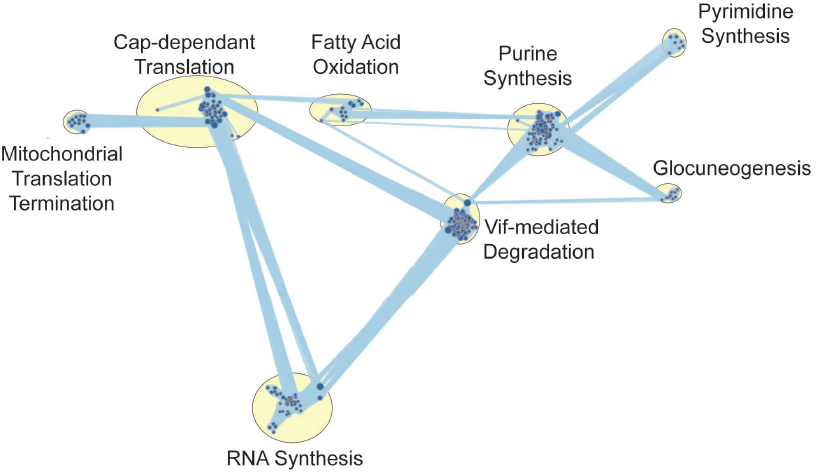
Pathway analysis reveals that euploid embryos significantly downregulate pathways involved in energy metabolism, transcription, and translation. When comparing euploid vs all aneuploid samples using GSEA, 156 gene sets were significantly enriched for downregulated genes at FDR<0.1. The size of the node reflects the number of genes in each gene set.

## DISCUSSION

In this work, we have shown that it is clinically feasible to explore the embryo’s transcriptome alongside its ploidy status from a single 4-6 cell trophectoderm biopsy. Notably, the addition of RNAseq did not interfere with the genomic analysis nor did it significantly alter the clinical quality control metrics (Table 3). A concordance of 100% makes us confident that this method could potentially be introduced into clinical use alongside the current work stream of PGT-A. The high level of concordance is a testament to the safety of this method, and the relatively large number of samples is a testament to the reproducibility of this method. This addition enhances the power of an existing tool to explore the potential of a specific embryo leading to a live birth, without the need for additional biopsies. Gaining insight into the transcriptomics of developing embryos could improve treatment choices as well as embryo selection and reduce time to pregnancy.

In the last 10 years, there has been a significant push to develop and apply single cell RNAseq methods (ISLAM *et al.* 2011; RAMSKOLD *et al.* 2012; SASAGAWA *et al.* 2013; TURCHINOVICH *et al.* 2014; CHAPMAN *et al.* 2015; HASHIMSHONY *et al.* 2016). Single cell-omics present many challenges, including limited starting material while at the same time generating a large amount of data. Recent advances in bioinformatics and the ability to integrate multiple layers of data (RNAseq, DNA methylation, histone modifications etc.) have driven the development of several methods for parallel sequencing of both mRNA and gDNA from the same cell or cell population. These methods rely on different ways of separating or differentiating between mRNA and gDNA (DEY *et al.* 2015; MACAULAY *et al.* 2015; REUTER *et al.* 2016).

We developed an approach that allows for the simultaneous sequencing of mRNA and gDNA from a low number of cells and which can be incorporated into the current clinical workflow used for ploidy testing. Furthermore, to facilitate the widespread adoption of simultaneous RNA sequencing in the clinic, the chosen kit/method must be easy to implement, sensitive, time-efficient, reproducible, and inexpensive. The method that best fit the above criteria for the low-input RNAseq arm of our project was a commercial development of SMART-seq (RAMSKOLD *et al.* 2012) (Takara Biosciences Inc.). By modifying the cell lysis step, we maintained the integrity of mRNA while additionally sequencing gDNA at a clinically acceptable quality (Figure 1).

Following our regular procedure of blastocyst biopsy for preimplantation genetic testing for chromosomal aberrations (PGT-A), we split the lysate and dedicated half to the transcriptome analysis. Upon splitting, all split samples (n=18) matched the ploidy of the unsplit (controls) “clinical samples” as shown in Table 3. With these results we can deduce a 100% concordance between gDNA sequencing data obtained following the current PGT-A process and that obtained following our integrated methodology. This data supports the safety and the reproducibility of our methodology.

When comparing between the transcriptomes of euploid and aneuploid embryos, we show that ploidy dramatically affects the transcriptome by PCA and hierarchical clustering (Figure 2). Previous studies correlating transcriptomics to ART outcomes in humans have done so without controlling for ploidy (JONES *et al.* 2008; KIRKEGAARD *et al.* 2015). Our current method not only employs NGS but is the first study to control for ploidy of the embryo when analyzing the full transcriptome. The importance of which is demonstrated by a 122-fold increase in the expression of *DDX19B* (involved in DNA repair) in T16 versus M16. This gene is located on the q-arm of chromosome 16 and the increased expression in T16 might simply be a result of the increased gene copy number due to trisomy and not necessarily biologically significant.

When comparing transcripts between euploid embryos and embryos with either T16 or M16, euploid embryos were enriched in several downregulated pathways involved in energy metabolism (fatty acid oxidation, gluconeogenesis, and mitochondrial translation termination) nucleotide synthesis, RNA transcription and protein translation (Figure 4). This is in agreement with the quiet embryo hypothesis, which states that embryo survival is increased by minimizing metabolism. These findings are further strengthened by several studies comparing metabolite usage in several commercial embryo culture media (BAUMANN *et al.* 2007; SWAIN 2015; GARDNER AND KELLEY 2017). *STOML2*, which is a key regulator of mitochondrial respiration, was upregulated in trisomy 16 embryos when compared with euploid embryos, further supporting the quiet embryo hypothesis. Interestingly, *FADS1* was upregulated in euploid when compared with aneuploid embryos, and is known to be involved in lipid metabolism and turnover, and has been previously shown to be abundant in elongating conceptus in sheep and to correlate with reproductive potential in cattle (CLEMENTE *et al.* 2011; BROOKS *et al.* 2016). Three genes important for implantation: *PGK1, PPAT*, and *EMP2*, which modulate angiogenesis and cell invasion, were differentially expressed between euploid and aneuploid embryos. Notably, previous studies on *EMP2* suggest that it may regulate implantation by orchestrating the surface expression of integrins and other membrane proteins (WADEHRA *et al.* 2005). *EMP2* was previously shown to be significantly reduced in both villous and extra-villous trophoblast populations in placentas of pregnancies with fetal intrauterine growth restriction (WILLIAMS *et al.* 2017).

When comparing our data to available data in the literature it should be noted that many factors can affect NGS results thus making it challenging to compare different datasets. Some of the factors that contribute to variability between different datasets are the method for obtaining the sample (either dissociating the embryos and picking up individual cells or utilizing TE biopsies) (VAN DEN BRINK *et al.* 2017), method of fertilization (intracytoplasmatic sperm injection or in vitro fertilization), culture conditions (open system vs closed culturing systems), fresh versus vitrified-thawed embryos and parental demographics (YAN *et al.* 2013; KIRKEGAARD *et al.* 2015; PETROPOULOS *et al.* 2016). However, these large data sets provide the groundwork/benchmark of the expected transcriptomic profile of TE cells at a specific time point.

The fields’ knowledge of the human embryo transcriptome has been restricted by limited access to human embryo research material and further complicated by the small amounts of available genetic material within one embryo. Furthermore, previous studies have predominantly relied on older techniques which have several limitations such as limited detection ability, high background levels and inherent bias. Finally, the most up to date studies have failed to control for euploidy status and haven’t incorporated their technique with what’s available today in the clinical setting. Our study has managed to improve on all those aspects. We have developed a method that has the potential to seamlessly incorporate into the clinical workflow without impacting the accuracy of aneuploidy detection by PGT-A. This allows us to reliably gain new insight into the transcriptome of embryos while controlling for ploidy, and the gives us the ability to correlate an embryo’s transcriptome to clinical outcome, such as implantation rate and live birth rate.

Once implemented into clinical practice, this novel multi-omic approach would help assess the impact different interventions have on the embryonal transcriptome. Furthermore, this novel tool might allow us to rank euploid embryos based on their potential to lead to live birth. In this study, we used our tool to assess the transcriptome of euploid and aneuploid embryos, and comparisons were made between the two cohorts with presumed different implantation and developmental potential. The fact that we could not correlate the findings with clinical outcomes (i.e. implantation rate, live birth rate etc.) is a limitation of this study and is the reason why we incorporated aneuploid embryos with presumed lower implantation and developmental potential in the design and validation of this tool.

Our novel approach applied in prospective studies will allow for interrogation of different clinical scenarios and treatments with the transcriptome of the developing blast and its potential to develop into a healthy baby. Our future research will focus on exploring in detail the validity of embryo transcriptome as a biomarker for embryo implantation and development. Differentially expressed genes between blastocysts that implant and ones that do not, while controlling for ploidy, would shed light on the pathways and genes involved in embryo implantation and enable further functional genome editing studies to enhance the knowledge on embryo implantation potential. We believe that uncovering the core of the transcriptomic map of embryo that leads to live birth will open the door for novel treatment options in ART including optimization of the culturing conditions to promote certain or critical metabolic pathways. Such an approach will potentially improve embryo selection for transfer as well as better define treatment factors that can influence the implantation potential of a given embryo, consequently improving the results in ART.

In conclusion, we have modified existing techniques to create a novel clinical tool. With declining costs of NGS and increased accessibility of that technology, along with advances in translational methods, we are closer than ever to being able to implement a multi omic approach when assessing the developmental potential of a single embryo. Further research is necessary to validate this technique and correlate it with clinical outcomes, with the goal of eventually incorporating it into the clinical setting.

## ACKNOWLEDGMENTS

The authors would like to thank all CReATe Fertility Centre embryologists, nurses, and staff for helping with data collection. We would also like to thank Ms. Kirah Hahn for helping with sample collection and cell manipulation in the early stages of this project.

## FIGURE LEGENDS

**Figure S1:** A) A representative image of a degenerated and late developing blastocyst deemed not suitable for transfer by an embryologist and used for lysis optimization. B) A representative image of a blastocyst being biopsied. Here the embryologist is gently aspirating 4-6 cells and removing this group of cells from the rest of the blastocyst. C) A representative image of a previously frozen blastocyst prior to re-biopsy for this study.

**Figure S2:** Bioanalyzer 2100 traces for all biopsied cells lysed using our novel PGT^2^ method and cDNA synthesized using the standard SMART-seq kit. All samples yielded high quality, full length cDNA.

**Figure S3:** qPCR validation of NGS data using four selected trophectoderm specific markers. All targets, except for CDX2, had similar expression by both qPCR and NGS.

**Table 1:** Characteristics of patients included in the study.

**Table 2:** Optimization of lysis method – VeriSeq next generation sequencing results from chromosomal aberration analysis and sequencing quality metrics

**Table 3:** VeriSeq next generation sequencing results from chromosomal aberration analysis and sequencing quality metrics from simultaneous RNA/DNA sequencing using the novel PGT^2^ method

